# Novel sampling strategies and a coarse-grained score function for docking homomers, flexible heteromers, and oligosaccharides using Rosetta in CAPRI Rounds 37–45

**DOI:** 10.1101/749317

**Authors:** Shourya S. Roy Burman, Morgan L. Nance, Jeliazko R. Jeliazkov, Jason W. Labonte, Joseph H. Lubin, Naireeta Biswas, Jeffrey J. Gray

**Affiliations:** Department of Chemical and Biomolecular Engineering, Johns Hopkins University, Baltimore, Maryland, USA; Department of Cancer Biology, Dana-Farber Cancer Institute, Boston, MA, USA; Department of Biological Chemistry and Molecular Pharmacology, Harvard Medical School, Boston, MA, USA; Program in Molecular Biophysics, Johns Hopkins University, Baltimore, Maryland, USA; Department of Biochemistry, University of Zurich, Zurich, Switzerland; Department of Chemistry, Franklin & Marshall College, Lancaster, PA, USA; Department of Chemistry and Chemical Biology, Rutgers, The State University of New Jersey, Piscataway, NJ, USA; FS-SCS, Deutsches Elektronen-Synchrotron, Hamburg, Germany; Institute for NanoBioTechnology, Johns Hopkins University, Baltimore, Maryland, USA; Sidney Kimmel Comprehensive Cancer Center, Johns Hopkins School of Medicine, Baltimore, Maryland, USA

## Abstract

CAPRI Rounds 37 through 45 introduced larger complexes, new macromolecules, and multi-stage assemblies. For these rounds, we used and expanded docking methods in Rosetta to model 23 target complexes. We successfully predicted 14 target complexes and recognized and refined near-native models generated by other groups for two further targets. Notably, for targets T110 and T136, we achieved the closest prediction of any CAPRI participant. We created several innovative approaches during these rounds. Since Round 39 (target 122), we have used the new RosettaDock 4.0, which has a revamped coarse-grained energy function and the ability to perform conformer selection during docking with hundreds of pre-generated protein backbones. Ten of the complexes had some degree of symmetry in their interactions, so we tested Rosetta SymDock, realized its shortcomings, and developed the next-generation symmetric docking protocol, SymDock2, which includes docking of multiple backbones and induced-fit refinement. Since the last CAPRI assessment, we also developed methods for modeling and designing carbohydrates in Rosetta, and we used them to successfully model oligosaccharide–protein complexes in Round 41. While the results were broadly encouraging, they also highlighted the pressing need to invest in (1) flexible docking algorithms with the ability to model loop and linker motions and in (2) new sampling and scoring methods for oligosaccharide–protein interactions.

## Introduction

With the explosion in genomic data availability and the ever-increasing accuracy of *in silico* protein folding methods, the ability to computationally model protein assemblies has taken center-stage. Protein-docking methods provide a rapid way to model assemblies, and hence, their progress has been a key focus of computational biophysics. Over the years, various approaches have been developed, each with a different scope and ability to integrate experimental data. Since 2001, a community-wide blind experiment, Critical Assessment of PRediction of Interactions (CAPRI), has been used to assess the state-of-the-art in computational macromolecular docking.^1^ Participating groups predict the structure of a complex given the sequences of the constituent proteins, stoichiometry of association, and, in case of homomeric complexes, the point symmetry. Based on their resemblance to the unpublished, experimentally determined structure, the accuracy of the predictions is ranked. With every new round, the organizers add to the complexity of the modeling challenge by introducing more intricate complexes and non-protein macromolecules.

Our group has continuously evaluated our docking algorithm, RosettaDock^2^ in CAPRI, leading to advances such as docking antibodies with loop flexibility,^3^ varying protonation states while docking,^4^ and interspersing conformational selection with docking.^5^ Previous rounds of CAPRI necessitated the creation of protocols for flexible protein assembly and oligosaccharide–protein docking.^6,7^ During this latest period with rounds 37 through 45, we developed RosettaDock 4.0 to model flexible proteins^8^ and refined GlycanDock to predict oligosaccharide–protein interactions.^9^ Round 37 was a joint experiment between CAPRI and the Critical Assessment of Structure Prediction (CASP) in which preliminary monomer models submitted by CASP12 participants were provided to the CAPRI participants for docking.^10^ This round comprised nine symmetric homomers. Based on our performance on these homomers, we developed a new symmetric docking algorithm called Rosetta SymDock2.^11^ Rounds 39 and 42 required global docking while predicting the conformation of long, flexible loops. These targets gave us an opportunity to add functionality to dock single-chain camelid antibodies in our antibody docking protocol, SnugDock.^3^

In this article, we examine the challenge of modeling with the available information, discuss the methodology we used, compare our CAPRI submissions to models made using new techniques, and suggest improvements to improve modeling accuracy.

## Methods and Results

We predicted the structures of 23 complexes in CAPRI rounds 37 through 45. We achieved 1 high-quality, 6 medium-quality and 7 acceptable predictions. Table I summarizes successfully modeled targets and Table II summarizes our failed attempts.

**Table I:**
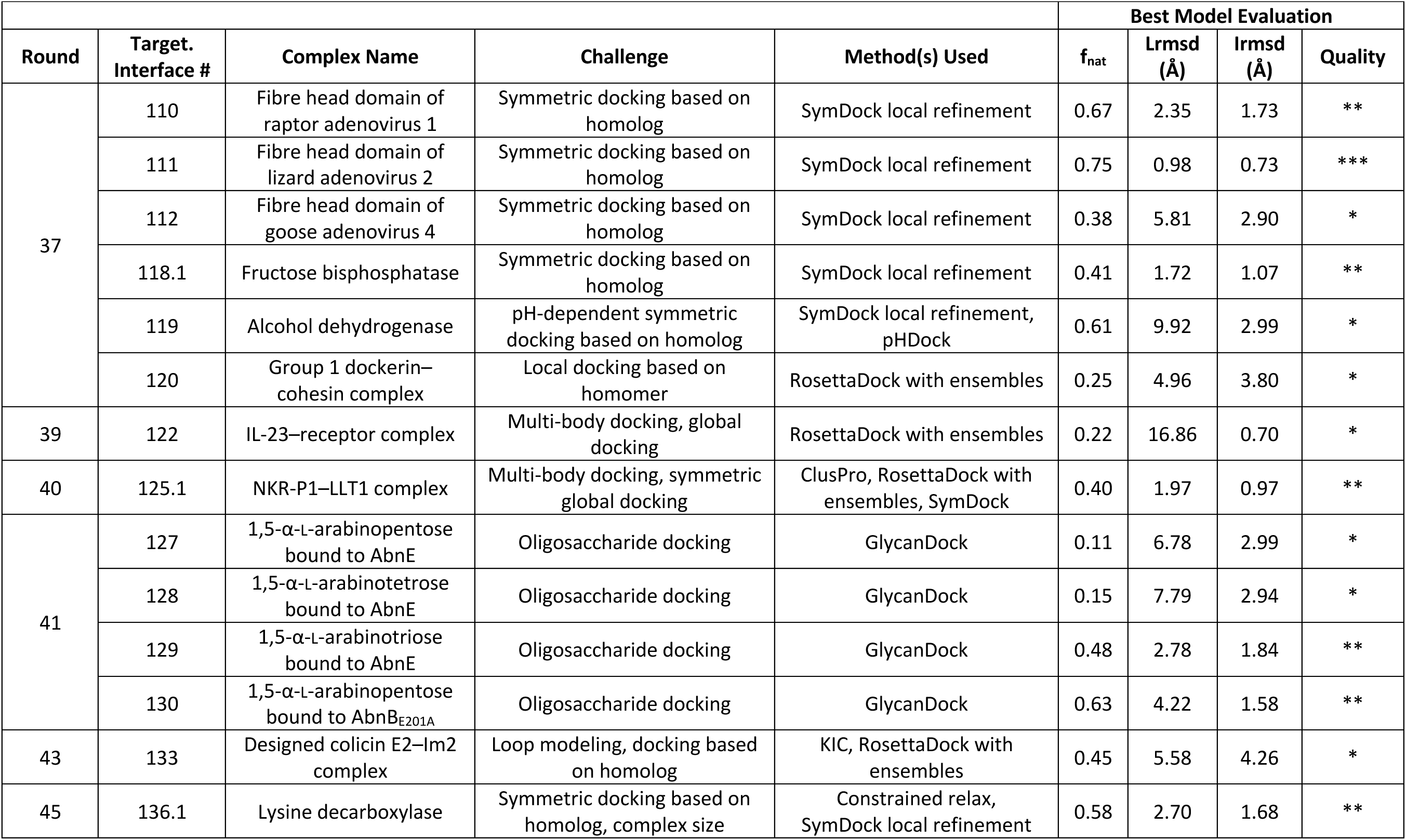
Summary of targets successfully modeled. The table lists the round, target number, name of the complex, the nature of the challenge, the methods used to model the complex, and the evaluation metrics for the best model that we submitted. The metrics are f_nat_: the fraction of native contacts recovered, Lrmsd: root-mean-square-deviation of the backbone atoms from the native ligand after superimposing the receptor, Irmsd: root-mean-square-deviation of the backbone atoms of the interface after superposition to the bound interface, and quality: high-quality (***), medium-quality (**), acceptable (*), or incorrect (-) as evaluated by the CAPRI organizers.

| Round | Target.<br>Interface # | Complex Name | Challenge | Method(s) Used | Best Model Evaluation |  |  |  |
| --- | --- | --- | --- | --- | --- | --- | --- | --- |
| | | | | | $f_{\text{nat}}$ | Lrmsd<br>(Å) | Irmsd<br>(Å) | Quality |
| 37 | 110 | Fibre head domain of raptor adenovirus 1 | Symmetric docking based on homolog | SymDock local refinement | 0.67 | 2.35 | 1.73 | ** |
|  | 111 | Fibre head domain of lizard adenovirus 2 | Symmetric docking based on homolog | SymDock local refinement | 0.75 | 0.98 | 0.73 | *** |
|  | 112 | Fibre head domain of goose adenovirus 4 | Symmetric docking based on homolog | SymDock local refinement | 0.38 | 5.81 | 2.90 | * |
|  | 118.1 | Fructose biphosphatase | Symmetric docking based on homolog | SymDock local refinement | 0.41 | 1.72 | 1.07 | ** |
|  | 119 | Alcohol dehydrogenase | pH-dependent symmetric docking based on homolog | SymDock local refinement, pHDock | 0.61 | 9.92 | 2.99 | * |
|  | 120 | Group 1 dockerin-cohesin complex | Local docking based on homomer | RosettaDock with ensembles | 0.25 | 4.96 | 3.80 | * |
| 39 | 122 | IL-23-receptor complex | Multi-body docking, global docking | RosettaDock with ensembles | 0.22 | 16.86 | 0.70 | * |
| 40 | 125.1 | NKR-P1-LLT1 complex | Multi-body docking, symmetric global docking | ClusPro, RosettaDock with ensembles, SymDock | 0.40 | 1.97 | 0.97 | ** |
| 41 | 127 | 1,5-α-L-arabinopentose bound to AbnE | Oligosaccharide docking | GlycanDock | 0.11 | 6.78 | 2.99 | * |
| | 128 | 1,5-α-L-arabinotetrose bound to AbnE | Oligosaccharide docking | GlycanDock | 0.15 | 7.79 | 2.94 | * |
| | 129 | 1,5-α-L-arabinotriose bound to AbnE | Oligosaccharide docking | GlycanDock | 0.48 | 2.78 | 1.84 | ** |
| | 130 | 1,5-α-L-arabinopentose bound to AbnB<sub>E201A</sub> | Oligosaccharide docking | GlycanDock | 0.63 | 4.22 | 1.58 | ** |
| 43 | 133 | Designed colicin E2-Im2 complex | Loop modeling, docking based on homolog | KIC, RosettaDock with ensembles | 0.45 | 5.58 | 4.26 | * |
| 45 | 136.1 | Lysine decarboxylase | Symmetric docking based on homolog, complex size | Constrained relax, SymDock local refinement | 0.58 | 2.70 | 1.68 | ** |

**Table II:** Summary of targets modeled incorrectly. ‘Best in CAPRI’ corresponds to the model evaluation of the best model submitted by all CAPRI predictors (not scorers). The description of the other headers is the same as in Table I.

| Round | Target.<br>Interface # | Complex Name | Challenge | Method(s) Used | Best Model Evaluation |  |  |  |
| --- | --- | --- | --- | --- | --- | --- | --- | --- |
| | | | | | $f_{\text{nat}}$ | Lrmsd (Å) | Irmsd (Å) | Best in CAPRI |
| 37 | 113 | CDI toxin–immunity complex | Flexible loop, global docking | ClusPro, RosettaDock with ensembles | 0.06 | 18.33 | 11.43 | * |
|  | 114 | Ljungan virus protein | Low-quality monomer, symm. global docking | SymDock with ensembles | 0.01 | 47.75 | 10.83 | - |
|  | 116 | Bifunctional histidine kinase | Variable domain linker, symmetric global docking | SymDock with ensembles | 0.09 | 36.02 | 11.86 | - |
|  | 117.2 | Pins–Insc complex | Partial monomer unfolding, multi-body docking | RosettaDock with ensembles, SymDock | 0.00 | 80.39 | 36.13 | - |
| 39 | 123 | PorM<sub>N-term</sub>–nb130 complex | Loop flexibility, global docking | RosettaAntibody, SnugDock, ClusPro | 0.16 | 18.53 | 5.06 | * |
|  | 124.2 | PorM<sub>C-term</sub>–nb02 complex | Loop flexibility, global docking | RosettaAntibody, SnugDock, ClusPro | 0.00 | 27.42 | 14.15 | - |
| 41 | 126<sup>a</sup> | 1,5-α-L-arabinohexose bound to AbnE | Oligosaccharide docking | GlycanDock | 0.00 | 11.24 | 4.27 | ** |
| 42 | 131 | CEACAM1–HopQ1 complex | Loop flexibility, global docking | CCD + fragments, RosettaDock with ensembles | 0.04 | 39.28 | 11.61 | * |
|  | 132<sup>a</sup> | CEACAM1–HopQ2 complex | Loop flexibility, global docking | CCD + fragments, RosettaDock with ensembles | 0.00 | 34.79 | 10.42 | ** |

<sup>a</sup> During the scoring round, we refined and scored models of other predictors to obtain acceptable structures.

### Homomer docking successes

#### Targets 110–112: Viral fiber head domains

The first three targets of round 37 were homo-trimeric fiber head domains from different viruses. Target 110 (T110) was the fiber head domain of raptor adenovirus 1, T111 was that of lizard adenovirus 2, and T112 was that of goose adenovirus 4, and the task was to predict the trimer quaternary structure.

Before we docked the models, the organizers provided us with the initial monomer models submitted by CASP12 participants. First, we relaxed all the models using Rosetta FastRelax,^12^ clustered those with similar backbone conformations, and chose between one and four monomer backbones for docking. For T110, all homologous proteins had less than 40% sequence coverage and identity. Importantly, all these distant homologs lacked a beta-hairpin predicted by CASP monomer models (residues 358 to 373) present in T110. T111 had a homolog with 94% sequence coverage and 50% identity, which strongly suggested that the native structure would resemble snake adenovirus 1 fiber head (PDB ID: 4D0V)^13^. For T112, the avian adenovirus CELO fiber head (2IUM)^14^ came the closest with 59% coverage and 27% identity. For T110, due to the aforementioned beta-hairpin, we performed symmetric global docking simulations with different monomer conformations, with and without the beta-hairpin, using Rosetta SymDock.^15^ For initial subunit placement for T111 and T112, we used subunit arrangements derived from their respective homologs and refined the complexes using fixed-backbone refinement of SymDock. Between 10,000 and 50,000 models per monomer were generated for all three targets.

For T110, the crystal structure (PDB ID: 5FJL)^16^ is shown in in Figure 1A (gray). The native structure did indeed possess a beta-hairpin as predicted, which is highlighted in red. On the superposed complex, the root-mean-square deviation of the C_α_ atoms (RMSD_Cα_) of the predicted beta-hairpin was 1.4 Å from the native. Our best model (yellow) recovered 67% of the native contacts across the subunits and had a root-mean-square deviation of ligand backbone atoms (Lrmsd) of 2.35 Å and a root-mean-square deviation of interface backbone atoms (Irmsd) of 1.73 Å from the native, which were the lowest values among all of the models submitted by all of the groups. The presence of a close homolog made T111 an easy target with multiple groups, including us, predicting high-quality models. The crystal structure of the target is still unreleased, but we assume it to be similar to snake adenovirus 1 fiber head (4D0V). In Figure 1B, our best model is shown in orange superimposed on the gray crystal structure of the homolog. Conversely, the absence of close homologs made T112 difficult to model both in the monomeric state and in the trimeric state. No group achieved a medium- or high-quality prediction. The Lrmsd and Irmsd of our best model was 5.8 Å and 2.9 Å, respectively and hence, it was classified as acceptable. The crystal structure of the target is still unreleased and no close homolog is available for a visual comparison.

**Fig 1:**
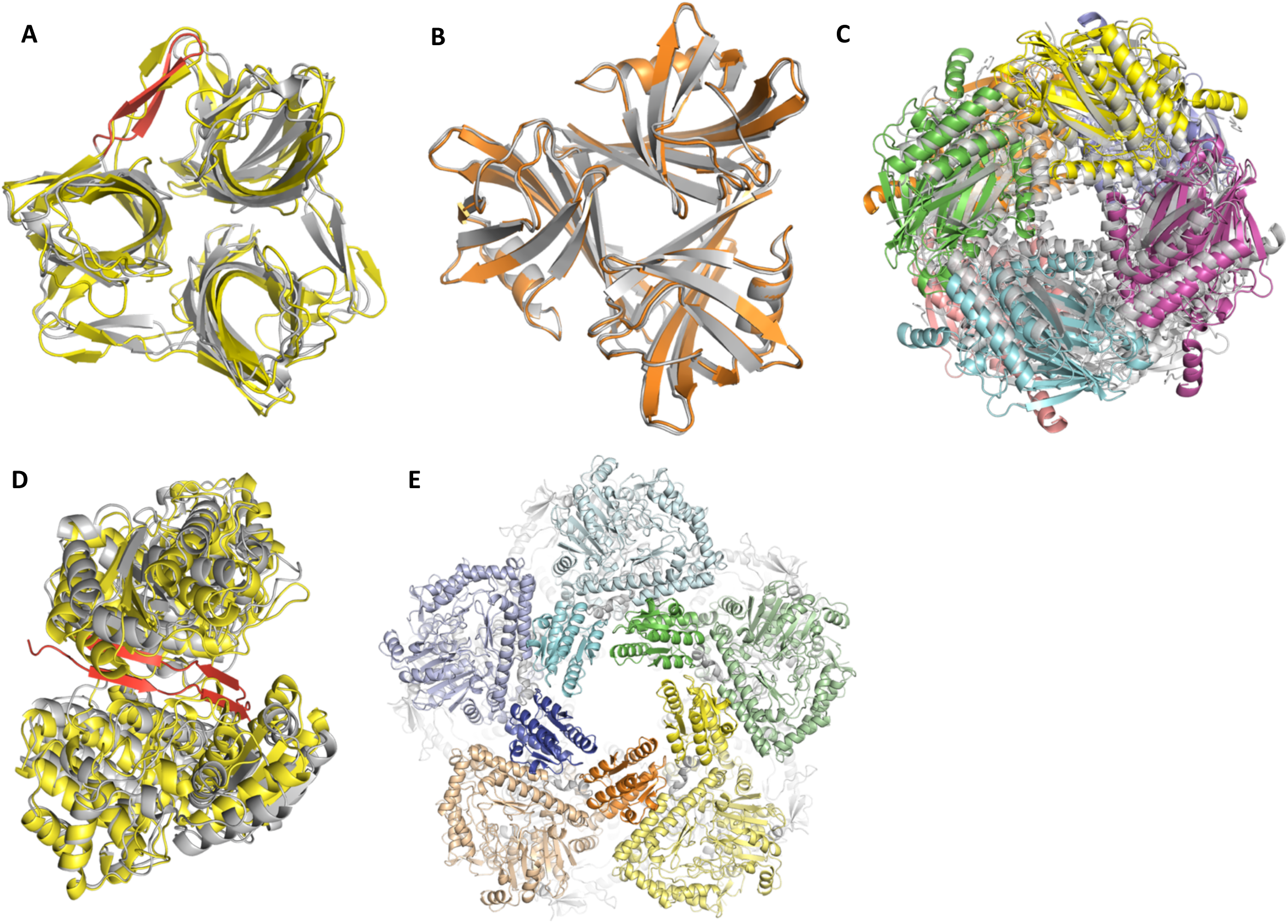
(A) **T110:** Our best model (** rating) of the fibre head domain of raptor adenovirus 1 (yellow) superimposed on the crystal structure (gray). Predicting and modeling the beta-hairpin in the native structure (red) was crucial to prediction success. The Lrmsd of 2.35 Å was the best among all groups. (B) **T111:** Our high-quality model (***) of the fibre head domain of lizard adenovirus 2 (orange) superimposed on the crystal structure of a close homolog (gray), snake adenovirus 1 fibre head. (C) **T118:** Our best model (**) of fructose 1,6-bisphosphatase (color) superimposed on the crystal structure of a close homolog (gray). (D) **T119:** Our best model (*) of alcohol dehydrogenase dimer (yellow) superimposed on the crystal structure (gray). The model is missing cross-beta interactions between the subunits at the interface (red). (E) **T136:** Our model (**) of LdcA decamer where the five subunits on top are displayed in different colors. For each subunit, the wing domains are highlighted in brighter hues. The Lrmsd of 2.44 Å was the best among all groups.

#### Target 118: Fructose bisphosphatase homo-octamer

T118 was a refinement challenge involving fructose 1,6-bisphosphatase from *Thermus thermophilus*. Although the organism is a hyperthermophile, we were not provided any temperature information about this target. A close homolog structure of fructose 1,6-bisphosphatase from a thermo-acidophilic archaeon, *Sulfolobus tokodaii* with 100% sequence coverage, 46% identity, and the same D4 symmetry (3R1M)^17^ was available.

We extracted symmetry information from the aforementioned homolog, arranged the monomer models and refined the complex using fixed-backbone refinement of SymDock. Figure 1C shows our best model in color and the crystal structure of the homolog in gray. The crystal structure of the target is yet to be released. The model recovered 41% of the native contacts with Lrmsd and Irmsd values of 1.7 Å and 1.0 Å, respectively, and hence, was classified as a medium-quality model.

#### Target 119: Archaeal halo-thermophilic alcohol dehydrogenase

T119 challenged us with atypical modeling conditions. The homo-dimeric protein, alcohol dehydrogenase, was from a halo-thermophilic archaeon expressed in a halo-mesophilic expression system. The behavior of this enzyme is pH dependent: in the pH range of 9.6–10.2, its oxidative reaction peaks, whereas at pH of 6.4, reduction reaction is dominant.^18^ We were asked to predict the structure of the complex at pH 10.

First, we relaxed and selected monomer models from CASP12 participants. The closest homologous homo-dimer that we found was alcohol dehydrogenase 2 from the bacteria *Zymomonas mobilis* (3OWO)^19^. The two subunits of the homolog had extensive cross-beta sheet interactions along the N-termini. The N-termini interaction served as a hinge, where a small error in the backbone would result in a drastically different rigid-body conformation. Unfortunately, this region of the target protein was predicted to be disordered and was different in all monomer models. As a result, we had to partly truncate the N-terminus. We followed a two-pronged approach to model this target: on the one hand, we explored the homo-dimeric conformational space using the standard SymDock protocol; on the other hand, we sampled different residue protonation states at pH 10 with a variant of RosettaDock called with Rosetta pHDock^4^. We produced 10,000 docking models with each method and chose the most symmetric proteins interacting at the N-terminus for pHDock.

Figure 1D shows our best model in yellow, the crystal structure in gray, and the N-terminus of the crystal structure in red. Despite missing key interactions at the N-terminus, we were able to predict the rough placement of the subunits correctly, and hence our best model was adjudged acceptable. The model predicted 61% of the native contacts with Lrmsd and Irmsd values of 9.9 Å and 3.0 Å, respectively. This large difference of RMSD values arises from the aforementioned hinge motion, where a small change in the N-terminus backbone leads to large changes globally. The best model across all groups was a medium-quality model.

#### Target 136: Lysine decarboxylase homo-decamer

T136 was the homo-decameric lysine decarboxylase, LdcA from *Pseudomonas aeruginosa*. Close homologs were available for the complex in the form of lysine decarboxylase, LdcI from *E. coli* (5FKZ) and arginine decarboxylase, AdiA (2VYC) from *Salmonella typhimurium*. The wing domain of the subunits was given to be significantly different to the homologs leading to different inter-subunit contacts. As the subunit arrangement was likely to be similar to its homologs, the challenge of this complex was to model the wing domain correctly within the confines of the D5 symmetry. Another issue was the sheer size of the protein: ten subunits each with 750 residues making extensive interfaces with other subunits makes it the largest CAPRI target to date, requiring high-memory workstations for docking.

We started by modeling the monomer using the online server, Robetta,^20^ which produced convergent conformations for the wing domain that were distinct from the homologs. Drawing on symmetry information from the homologs and arranging the subunits accordingly led to steric clashes at the wing domain. All fixed-backbone refinement efforts using SymDock failed to produce a plausible structure free of clashes. Instead, SymDock refinement resulted in an unrealistic inter-subunit distance by expanding the complex to relieve the clashes. We conjectured that the monomers needed to be relaxed in the context of the complex, and not independent of it. We achieved this by superimposing ten copies of each monomer model onto each of the subunits of the homolog, AdiA and then relaxing one of the monomers with the other nine copies present. On doing so and then docking the context-refined monomers, we were able to obtain structures where the wing domain readily fits into the given symmetry without steric clashes. Figure 1E shows our model for LdcA with the wing domains of five subunits highlighted in darker hues against lighter tints of the rest of the subunit. We predict that each wing domain contacts two neighboring wing domains and well as a neighboring chain. Our scoring model (based on our prediction model) was adjudged to be the highest quality for any group (medium quality overall) and recovered 60% of the native contacts with Lrmsd and Irmsd of 2.4 Å and 1.7 Å, respectively.

#### Post-hoc analysis of the performance of SymDock2 on CAPRI targets

We noticed a pattern of error whereby when starting from the symmetric arrangement of a homolog, Rosetta SymDock protocol would expand the overall size of the complex to relieve inter-chain clashes. This phenomenon progressively worsened for higher order symmetries. Given the same set of inputs, we tested (post-CAPRI) whether SymDock2^11^ improved the quality of the models for three of the complexes where this error was observed, *viz.* T110, T118, and T136.

For T110, the best model amongst the 10 top-scoring models had the same overall classification as our original submission, medium-quality. However, its inter-subunit distance of 23.7 Å was 2% smaller than the native (5FJL) structure’s distance of 24.2 Å. This reversed the trend of the best structure from SymDock, which had a 4% larger inter-subunit distance of 25.0 Å. The cause of this compression is not because the individual monomers are closer to the native; in fact, the RMSD_Cα_ of the monomers in SymDock was 1.1 Å compared to 1.2 Å after SymDock2. The tighter fit was achieved by subtle backbone changes during SymDock2’s flexible backbone refinement.

We observed a big improvement for T118, where most of SymDock2’s 10 top-scoring models were high-quality (using monomer model superposition to a close homolog, 3R1M, to approximate native). The best structure from SymDock2 recovered 73% of the “native” contacts while having a sub-angstrom Lrmsd. Had we submitted this structure during CAPRI, it would have been the best structure across all groups. Moreover, the inter-subunit distance of the SymDock2 model was 40.3 Å compared to 43.3 Å in the SymDock model and 39.0 Å in the homolog. Thus, the SymDock2 model expands by just 3% relative to the homolog and presumably recovers additional inter-chain contacts compared to the SymDock model, which expands by 11%. Supplementary Figure S1A highlights the similarity of the SymDock2 model to the “native” and Figure S1B compares the SymDock and SymDock2 models and the distance between monomers.

As we could not generate a feasible structure for T136 using SymDock, a direct comparison is not possible. Instead, first, we had relaxed the monomer in context of its partners and then used that monomer with SymDock. The inter-subunit distance for the best-scoring model was 62.2 Å, a 1% increase over the distance of 61.6 Å in a close homolog (2VYC). With SymDock2, we could generate models starting from the initial homology-modeled monomers with the best-scoring model having an inter-subunit distance of 63.1 Å, which was 2% more than the homolog. Thus, with better packing of interfaces due to flexible backbone refinement, SymDock2 resolves the problem of complex expansion in SymDock and thus, significantly outperforms SymDock on CAPRI targets.

### Homomer docking failures

#### Target 114: Ljungan virus protein

T114 was the homodimeric-protein 2A2 from Ljungan virus. The function of this protein is unknown. We were provided with monomer models from CASP12 participants. We relaxed the models and chose five top-scoring, distinct models for docking. We found no homologs from which to extract symmetry information, and hence, we performed a global search of C2 configuration space to generate 50,000 models for each monomer structure. None of the models submitted by us or any predictor was adjudged to be correct. As the experimental structure of this protein has not been released yet, we could not determine the reason(s) for failure.

#### Target 116: Bifunctional histidine kinase

T116 was two domains (Dhp and CA) of the homo-dimer, CckA of *Caulobacter crescentus*. We identified several homo-dimeric histidine kinase homologs with 97% or more sequence coverage and 25% or more identity, like those from *E. coli* (4GCZ)^21^ and *Geobacillus stearothermophilus* (3D36)^22^. Each homolog had a different relative orientation of the Dhp and CA domain equivalents, indicating that this target could only be successfully docked if the domains were correctly oriented in the monomers.

Models from CASP12 participants had a variety of different relative orientations of these two domains depending on the homolog template they chose. Using monomers with two different orientations, we generated 25,000 global docked models per orientation. Unfortunately, the relative orientations of the two CckA domains were very different from all available homologs as shown in Supplementary Figure S2B. Without a good monomer conformation, we, as well as all the other predictors, failed to dock the dimer correctly.

### Heteromer docking successes

#### Target 120: Group 1 dockerin–cohesin complex

In anaerobic bacteria, the cellulosome assembly digests plant fibers. The assembly of this complex involves the binding of different enzyme-borne dockerin proteins (Doc) to cohesin modules of the non-enzymatic protein, scaffoldin (Sca). As different groups of dockerins have significantly different cohesin-binding interfaces, they have different binding modes for every cohesin.^23^ Moreover, within the same species, each dockerin binds cohesins promiscuously and with different binding modes. Furthermore, single residue changes can affect binding, compounding the challenge of identifying the correct mode.^24^ T120 was a hetero-dimer of ScaB3 cohesin with Doc1a from *Ruminococcus flavefaciens*.

We started with homology models from CASP12 participants and relaxed them. Next, we searched for homologous complexes to create an initial placement of the monomers. A complex of group I dockerin and ScaB from *Acetivibrio cellulolyticus* (4UYQ)^25^ provided a starting point despite low homology with the individual proteins—the cohesin had 24% sequence coverage with 33% identity and the dockerin had 79% coverage with 37% identity. Starting from an initial structure where the monomers were aligned to the homologous complex, we docked the target. While docking the proteins, we used an ensemble of 10 relaxed monomer models for each partner to explore alternate backbone conformations.

Figure 2A shows our best model in green (cohesin) and blue (dockerin) against the crystal structure in gray. This model was adjudged to be acceptable; no other group submitted a higher-ranking model. The bulge in the crystal structure of cohesin (highlighted in red) was not present in any of the homology-modeled cohesins. This bulge changed the rigid-body conformation of the dockerin and resulted in the dockerin having an Lrmsd of 4.9 Å.

**Fig 2:**
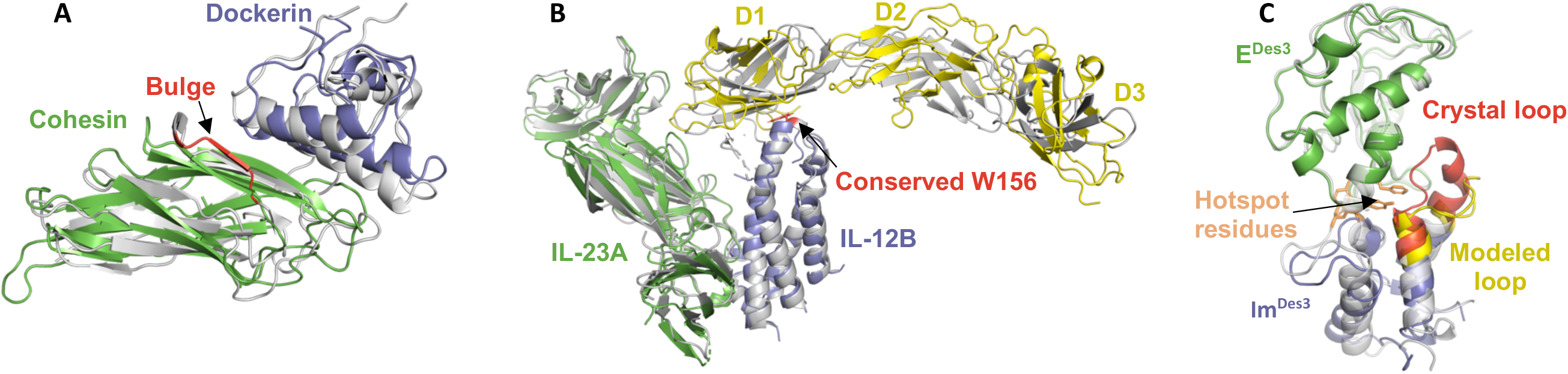
(A) **T120:** Our best model (*) of group I dockerin (blue)–cohesin (green) complex superimposed on the crystal structure (gray). A bulge in the cohesion (red) was not modeled correctly leading to a small error in the rigid body orientation of the dockerin. (B) **T122:** Our best model (*) of the complex of the two chains of IL-23, viz. IL-23A (green) and IL-12B (blue) with IL-23R (yellow) superimposed on the crystal structure (grey). The lightning rod interaction via the conserved Trp-156 (red) to IL-23R domain 1 (D1) was correctly predicted. IL-23R model had large errors in the relative orientation of the three domains. (C) **T133:** A 14^th^-ranked model (**) of Im^Des3^ (blue) bound to E^Des3^ (green) superimposed on the crystal structure of the complex (gray). The Im^Des3^ loop (red) is modeled inaccurately (yellow). This alters the rigid body orientation of Im^Des3^ by pushing it ‘down’. The hotspot resides across the interface in the wild-type (orange), *viz.* Y54-Y55 Im2 and F86 on E2 still interact in the model of the designed complex.

#### Target 122: Human IL-23–receptor complex

For T122, we were asked to model the interaction between IL-23 and its receptor, IL-23R. Several crystal structures of IL-23 were available in the Protein Data Bank.^26^ A disulfide bond held together its two subunits, IL-23A and IL-12B and hence, we expected their bound state to remain largely unchanged. We modeled the receptor, IL-23R using Modeller^27^ based on multiple sequence alignment of homologs with manual input on the alignment of loop regions. In addition, we also used models from Robetta,^20^ which used a different homolog as its template.

From the variety of models obtained, it was apparent that the receptor might have inter-domain flexibility between its three domains. This flexibility ruled out the possibility of global docking. A literature survey revealed that the binding site observed in other cytokine/cytokine receptor complexes in this family was likely used to bind IL-12Rβ1 (which was not the receptor chain we were modeling) and not IL-23R.^28^ Based on prior experimental experience on IL-23 interactions, a collaborator (Jamie Spangler) advised us that the interaction was likely between the D1 domain of IL-23R and IL-23 with the conserved Trp-156 on IL-12B serving as the ‘lightning rod’. Using this information, we obtained a starting state and locally docked the receptor against the cytokine heterodimer while constraining the conserved tryptophan residue to contact the receptor. This was the first target for which we used RosettaDock 4.0, and as a result we were able to efficiently dock 65 receptor backbone conformations to 56 cytokine backbone conformations.

Figure 2B shows our best model superimposed on the crystal structure (5MZV, in gray)^29^. The conserved tryptophan of IL-12B is highlighted in red. This model was able to capture the rough binding mode along with the tryptophan lightning rod interaction. With 22% of the native contacts recovered and an Irmsd of 0.7 Å, our model was adjudged acceptable. None of the models of IL-23R (yellow) had RMSD_Cα_ under 4.2 Å because of the different orientations of the three domains and as a result the Lrmsd of the model was 16.8 Å. The best model across all the CAPRI groups was a medium-quality model with 40% of the native contacts.

#### Target 125: NKR-P1–LLT1 hetero-hexamer

T125 was the complex between the extracellular domains of natural killer cell surface receptor, NKR-P1 and a cell surface ligand, LLT1. It presented a three-step docking challenge: first, a dimer of NKR-P1 had to be modeled, then the LLT1–NKR-P1 hetero-trimer complex had to be determined, and finally, two of these hetero-trimers had to be docked together to construct the hetero-hexamer.

For NKR-P1, we generated dimer models by symmetric docking of the monomer models of NKR-P1 obtained from Robetta. We chose seven dimer configurations for further docking. We then modeled the NKR-P1 dimer–LLT1 complex by global docking of the models using ClusPro followed by local refinement in Rosetta. A structure of LLT1 dimer was already available (4QKH); we used this as a reference to assemble the complex model. This step also filtered out trimer configurations that clashed with each other. Finally, we locally refined three candidate complexes, generating 5,000 models each.

Our best model captured 40% of the native contacts on the LLT1–NKR-P1 with an Irmsd of 0.969 Å and Lrmsd of 1.971 Å and was classified as medium-quality. On *post ex facto* analysis, the closest docked conformation had an RMSD_Cα_ of 4.7 Å from the crystal structure of NKR-P1 homodimer (5MGS). As a result, we, as well as other predictors, could not predict the full hexamer correctly.

#### Target 133: Colicin DNase–immunity protein complex

T133 was a colicin E2 DNase–Im2 complex designed to change partner specificity from the native complex. The crystal structure of the native colicin E2 DNase–Im2 was available (3U43).^30^ However, the organizers informed us that the mutations led to an altered binding mode. Therefore, the challenge of this target was recognizing changes in the binding mode brought about by the mutations. The designed colicin, E^Des3^ had mutations in 17 of the 132 positions while the immunity protein Im^Des3^ had 15 of its 85 positions mutated, most of which were situated in a loop. Three residues, identified as native-sequence hotspots for binding (Y54 and Y55 on Im2 plus F86 on E2),^30^ were not changed. After mutating and refining the structures of the mutant proteins, we explored different conformations of the Im^Des3^ loop with the mutations (residues 20–35) and closed the loop with kinematic closure.^31^ For E^Des3^, we obtained a variety of backbone conformations using Rosetta Backrub.^32^ We then docked an ensemble of E^Des3^ conformations with an ensemble of Im^Des3^ conformations while constraining the three hotspot residues to interact.

Figure 2C shows our best E^des3^ (green)–Im^Des3^ (blue) model (submitted as our 14^th^ model) superimposed with the crystal structure (gray) of E^des3^– Im^Des3^ complex (6ERE)^33^. Unfortunately, we predicted larger backbone changes (yellow) than were actually a part of the design (red). The hotspot residues (orange sticks) interact as predicted. This medium-quality model predicts 42% of native contacts with 2.2 Å Irmsd and 4.1 Å Lrmsd. We only submitted acceptable structures in our top ten.

### Heteromer docking failures

#### Target 113: Contact-dependent toxin–immunity protein complex

In T113, we were asked to model the interaction between the C-terminal domain of the toxin, CdiA-CT, and its cognate immunity protein, CdiI2 from *Cupriavidus taiwanensis*. We started with monomer models from the CASP12 predictors. We observed variability in the CdiA-CT models and consequently chose nine that had convergent secondary structure signatures for further modeling. There was less variability in CdiI2 models, but for the eleven-residue N-terminal tail. We chose three models with significantly different tail conformations from each other to hedge our bets. As we could not find a homologous complex, we searched the global conformational space using ClusPro^34^ and chose the binding mode compatible with most of the monomer conformations. Restricting our search to the local space around this mode, we then docked the ensemble of nine CdiA-CT backbone with three CdiI2 backbones to generate 15,000 models. As we could not predict this tail conformation correctly in the model, we predicted the rigid body conformation of CdiA-CT incorrectly (see Supplementary Fig S2A). The best model across all groups was classified as acceptable.

#### Target 117: Pins–Insc tetramer

T117 was the tetrameric complex of two molecules of Pins, a cell polarity determining protein and two molecules of Insc, an adapter protein. A structure of the Pins monomer was available (3SF4). For the structure of Insc, we relied on models from CASP12 participants. Owing to the absence of close homologs, we obtained a variety of different models, which we then relaxed and clustered by similarity. The models that clustered most tightly still had a variety of conformations of the first 35 (N-terminal) residues, which we consequently truncated. Based on literature,^35^ we decided to construct a homo-dimer of hetero-dimer model, where we first docked the Insc to Pins, generating 50,000 models, then selected an ensemble of 15 distinct top-scoring dimers, and finally we symmetrically docked the dimers, starting from four distinct orientations, and generating 50,000 models for each orientation.

The crystal structure of the complex (5A7D, see Supplementary Fig S2C) is a homo-dimer of two hetero-dimers as we predicted, but is not symmetric. The primary contacts of Insc in each hetero-dimer unit occur in the thirty-residue unfolded N-terminal peptide, Insc^PEPT^. As a result, there is a large amount of conformational flexibility in the hetero-dimer subcomplex with the two subcomplexes in the crystal structure having significantly different conformations. We had truncated this peptide and hence could not model either the hetero-subcomplex or the whole complex correctly. This was arguably the hardest challenge of round 37 because it involved not only multi-body docking, but also predicting the interactions of an unfolded peptide stretch with large conformational flexibility. No CAPRI team submitted an acceptable or better model.

#### Targets 123 & 124: PorM–camelid nanobody complex

T123 and T124 comprised the N- and C-terminal domains (respectively) of PorM, a periplasmic member of the type IX secretion system found in *Porphyromonas gingivalis*, in complex with nanobody chaperones. In their presence, the N-terminal domain crystallized as a monomer whereas the C-terminal domain crystallized as a dimer. Generally, nanobodies recognize antigens primarily by interactions in three variable loops called H1, H2 and H3. The H3 loop is the longest and most flexible loop, and as a result it is the primary determinant of binding. Thus, we modeled the constant core of the nanobody and the H1 and H2 loops from available homologs and then we generated 1,000 models with different H3 loop conformations using RosettaAntibody.^36,37^In T123, PorM_N-term_ in complex with nb02, the H3 loop was 12 residues long, whereas in T124, PorM_C-term_ dimer in complex with nb130, the H3 was 21 residues long.

For T123, we obtained PorM_N-term_ models from Robetta. For T124, we obtained PorM_C-term_ monomer conformations from Robetta and docked them together symmetrically to attain a dimer configuration. In both cases, no homologs were available as templates and hence the monomers were modeled *de novo* from sequence, which was a source of error. Using the lowest-scoring PorM models, we searched for suitable nanobody-binding regions by global docking using ClusPro. We then refined the distinct binding modes obtained from ClusPro while simultaneously sampling various nanobody variable loop conformations using SnugDock,^3,36^ a variant of RosettaDock specialized for docking antibodies.

For T123, the PorM_N-term_–nb02 complex, we did not produce an acceptable model or better. Since the structure is not yet released, we cannot analyze the reason for our failure. Only one acceptable solution was submitted across all participants. T124, the PorM_C-term_ dimer–nb130, complex involved multiple challenges: modeling monomers without templates, predicting their dimeric form and then docking the nanobody correctly (see Supplementary Figure S3A). All our PorM_C-term_ monomeric models were incorrect. Although we were able to model the nb02 H3 loop within 2.4 Å RMSD_Cα_, all our complex models had an Lrmsd of more than 18 Å. As a result, we, as well as all the other participants, failed to model this complex correctly.

#### Targets 131 & 132: CEACAM1–HopQ complex

T131 and T132 were the complexes of *Helicobacter pylori* adhesins HopQ1 and HopQ2, respectively, bound to the N-terminal domain of cell adhesion molecule CEACAM1. Multiple structures were available for the N-terminal domain of CEACAM1 (2GK2, 4QXW, 4WHD, and 5DZL).^38,39^ For T131, the structure of HopQ1 was available with four loops missing at the putative binding interface (5LP2).^40^ Using this structure as a template, complete models were obtained from Robetta. Robetta produced different conformations for the two longest loops (residues 135–148 and 245–255), which suggested potential flexibility. A mutation study indicated that residues Y34 and I91 of CEACAM1 are essential for HopQ binding.^41^ The authors of the study also conjectured that the first long loop of HopQ1 is involved in binding CEACAMs. We modeled the missing loop using fragment insertion and closed the loop with cyclic coordinate descent.^42^ Using an ensemble of 200 different loop conformations for the first HopQ1 loop and constraints to ensure CEACAM1 Y34 and I91 contact HopQ1, we generated 10,000 models each from two different starting states. For T132, we modeled the structure of the HopQ2 monomer based on its homology to HopQ1 using Rosetta Remodel.^43^ We followed a similar protocol for HopQ1 loop conformation sampling (for a slightly shorter loop of residues 135–144) and docking.

In both the cases, our loop modeling methods failed to provide the necessary bound conformation, often producing extended loops, instead of the compact structure in the crystal. As a result, the rigid-body orientation of CEACAM1 was completely incorrect. The two CEACAM1 residues predicted to be at the interface were indeed found to be there and are shown as salmon sticks in Supplementary Figure S3B.

While we did not predict the structure correctly, we did successfully refine and score structures submitted by another group. Our best refined model was classified as acceptable with 27% of native contacts predicted, Lrmsd and Irmsd of 11.8 Å and 3.2 Å, respectively. This demonstrates that the *REF2015* score function^44^ can recognize the near-native structure. Therefore, the outstanding challenge is to sample the conformation *de novo*.

### Protein–Oligosaccharide docking

#### Targets 126–130: Arabino-oligosaccharide binding to proteins

In round 41 of CAPRI, we modeled the interaction between arabino-oligosaccharide ligands of different lengths and the arabinose sensor, AbnE, or the arabinanase, AbnB — two important components of the L-arabinan-utilization system of *Geobacillus stearothermophilus*. Specifically, T126–129 challenged us with the docking of 1,5-α-L-arabinohexose (A6) through 1,5-α-L-arabinotriose (A3), respectively, to AbnE. T130 involved the docking of A5 to a catalytic mutant (E201A) of AbnB.

We modeled AbnE from homologs with 95% or more sequence coverage and 25% or more identity using Modeller^27^ and relaxed the models in Rosetta.^45^ Additionally, we obtained models from the Robetta server.^20^ One of the homologs that we used to model the target, the maltose-binding protein GacH from *Streptomyces glaucescens*, exists in two conformations: an unliganded open conformation and a closed, ligand-bound conformation.^46^ From all the aforementioned protein models, we used the conformation closest to the ligand-bound GacH conformation to model T126–129. As the chemical description of arabinose was absent in Rosetta, we programmed the required geometry, partial charge, and chemical connectivity information to model arabinose ligands with the RosettaCarbohydrate framework.^9^ To obtain a starting structure, we superimposed the AbnE model and A4 onto maltotetraose-bound GacH (3K00)^46^, changing the backbone torsion angles of A4 to best align with maltotetraose. For A5 and A6, we added arabinose units to the non-reducing end of the ligand. For A3, we removed an arabinose unit from the non-reducing end.

To simultaneously dock the glyco-ligands and explore their backbone conformations, we used the new GlycanDock protocol in Rosetta.^9^ In this protocol, the glyco-ligand undergoes small backbone motions along with rigid-body moves to dock into a protein cavity. These perturbations are alternated with side-chain repacking and energy minimization in torsion space for the residues at the protein–glycan interface. For each target, we obtained 15,000 initial docked models without any constraints to relieve clashes and to broadly sample the rigid-body conformational space. From the models where the ligands moved less than 5 Å RMSD from the starting structure, *i.e.*, those that stayed in the binding pocket of AbnE, we selected the one with the lowest interaction energy as the starting model for the final simulation. For the final docking simulation, we added constraints to hold the glyco-ligands within the putative binding pocket of AbnE and generated another 15,000 models. The range of conformations explored by A6 in T126 is exemplified in Figure 3A. We predicted medium quality models for T129 and acceptable-quality models for T127 and T128. During scoring, we recognized acceptable models for T126–T129, thus validating the score function.

**Fig 3:**
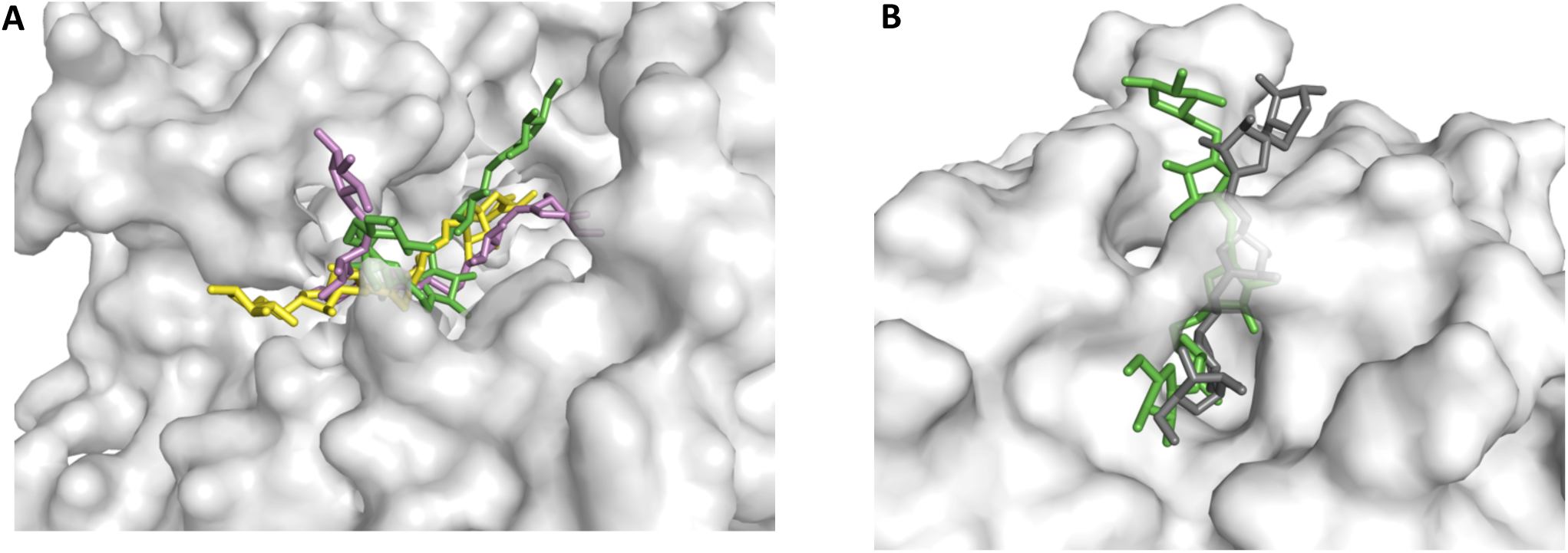
(A) **T126:** Range of ligand conformations sampled by 1,5-α-L-arabinohexose (green/yellow/pink) in the binding groove of AbnE (gray). (B) **T130:** The best predicted conformation of 1,5-α-L-arabinopentose (green) on AbnB_E201A_ (gray), with *f*_nat_ = 63%. The crystal structure (dark gray) is shown for comparison.

For T130, the crystal structure of A3 bound to the E201A mutant of the glycosidase AbnB was already available (3D5Z). The active site of this enzyme is a long groove with a bridge connecting the brinks under which the ligand can slide (see Figure 3B). The enzyme cleaves glycosidic linkages indiscriminately^47^ as the groove offers no steric obstruction at either end to hold the substrate in place. Consequently, although a structure was available with A3, we could not *a priori* predict how the A5 ligand would position itself. Extending the A3 in either direction provided us with starting coordinates for three unique starting states. We generated 10,000 docked models each from the three starting states using GlycanDock while constraining the A5 ligand to the active site groove. Figure 3B shows the best predicted conformation of A5 (in green) superimposed on the crystal structure of AbnB_E201A_ (light gray)–A5 (dark gray) complex (6F1G). The structure is a medium-quality model with 63% of the contacts being recovered with an Irmsd of 1.58 Å and an Lrmsd of 4.22 Å.

#### Post-hoc analysis of GlycanDock sampling and scoring

Compared to the crystal structure (6F1G), the three A5 structures we used as input for the T130 prediction round had a high fraction of native contacts (0.267, 0.367, and 0.633, respectively). We examined the failure of Rosetta GlycanDock to produce a high-quality structure despite having favorable starting states by investigating the Rosetta scoring function and an updated sampling algorithm in the version of GlycanDock under development.

To test scoring, we refined the crystal structure to generate 50 models. All models had sub-Angstrom RMSDs (purple triangles in Supplementary Figure S4) with interaction scores more favorable than those of the submitted models; the lowest-scoring model had a score of −24.9 units for the crystal refinement versus −13.9 units for the submitted models. This result suggests that, had it been sampled, the Rosetta scoring function would have correctly identified a near-native structure.

Despite having favorable starting conformations and a score function that discriminates near-native models, the previous version of GlycanDock failed to sample native-like states. The primary reason was that the first step of the algorithm randomized glycan backbone torsions as well as the rotation of the glycan about the protein, both of which disrupted favorable starting structures. As a result, this initial perturbation is not as extensive in the updated version.

To diagnose our current limitations, we tested two aspects of sampling—the rigid-body orientation sampling and the glycan backbone conformation sampling—individually at first and then, simultaneously. To remove any bias from the protein backbone conformation in the crystal, we used the protein backbone that we used for the prediction round. As a best-case scenario, we aligned the crystal structure of the A5 glycan in the protein groove as observed in the crystal structure. We generated 500 models by perturbing the glycan conformation (0.5 Å translation, 7.5° rotation, and backbone torsion perturbation between ± 12.5°) and then refining it. As expected, we successfully generated and discriminated near-native decoys even when starting with structures with an average Lrmsd of 1.98 Å (Supplementary Figure S4A). Next, we examined rigid-body sampling by moving the A5 crystal structure away from the binding pocket by assigning it *de novo* coordinates. Starting with this orientation and employing the same protocol, we generated high-quality models with RMSDs similar to those observed in crystal structure refinement (Supplementary Figure S4B). To examine glycan backbone sampling, we aligned the three input A5 structures used for the prediction round in the protein groove as in the crystal structure and generated 1000 docked models for each input. We were unable to obtain any high-quality models; all models had Lrmsd greater than 2 Å (Supplementary Figure S4C). Finally, we tested both rigid-body and backbone sampling simultaneously by placing the three input A5 structures used for the prediction outside the binding groove and then docking. We achieved similar results as in the previous case (Supplementary Figure S4D). These results suggest GlycanDock adequately samples rigid-body orientations but fails to do so for glycan backbone conformation. Thus, the key to successfully dock glycans lies in sampling relevant glycan backbone torsion.

## Discussion

Previous rounds of CAPRI led to the development of niche protocols like SnugDock^3^ to model antibody–antigen binding and pHDock^4^ to dynamically sample residue protonation states while docking.^6^ In rounds 37–45, we utilized these specialized methods while also encountering challenges that require overhauls of the core methodology for general problems such as global docking with flexibility, global docking of symmetric homomers, and oligosaccharide–protein docking. We modeled backbone flexibility by incorporating a pre-generated ensemble of backbone conformations during docking. With RosettaDock 4.0,^8^ we sampled over fifty conformations for each partner to successfully model T122. Despite having an efficient backbone sampling algorithm, we failed to model T131 and T132 due to the absence of conformations where the interacting loops were in near-bound conformation. These failures highlight the need for ensemble generation methods that sample loop conformations broadly.

As many of the targets were symmetric homomers with varying degrees of homology to existing structures, we were able to thoroughly assess the Rosetta SymDock protocol. When homologs were present, we could borrow the symmetric arrangement from the homolog as a template, as we did to successfully model targets 110, 111, 112, 118, and 119. However, even in those cases, the proximity of the monomer backbone to the template monomer backbone determined the overall quality of the models. For example, the monomer model of T111 had a 0.8 Å RMSD_Cα_ from the template and was our only prediction to be classified as high-quality. While one would expect that the more closely related a template is, the better the model will be, we noticed a systematic pattern of error in tightly-packed, higher order complexes. The method of induced fit successfully used for T136 inspired the flexible-backbone refinement strategy of the new symmetric docking protocol, Rosetta SymDock2.^11^

For only the second time in CAPRI, we encountered oligosaccharide–protein complexes. Five targets in round 41 gave us an opportunity to work with the recently-developed RosettaCarbohydrate framework,^9^ especially the GlycanDock application therein. Oligosaccharides have many more degrees of freedom than peptides, often featuring an additional mobile backbone torsion angle, multiple mobile side chains, and sometimes flexible rings. GlycanDock samples these mobile dihedrals while performing rigid-body transformations to place the oligosaccharide in a binding pocket and simultaneously repacking the side chains of contacting protein residues. We recognized deficiencies in the sampling of GlycanDock, and ongoing developments focus on optimizing its conformation sampling capabilities for a variety of glyco-ligands. Certain glycosidic linkages have been observed to populate limited regions of torsion space and for these linkages, glycosidic torsion angle preferences and crystal structure-based statistics have been calculated and collected.^48,49^ For the arabinose–arabinose linkages present in T130, linkage torsional statistics have not yet been collected, nor have the linkage-conformation energies been calculated. These data, when incorporated into glycan docking algorithms, could narrow the search space in glycan conformation sampling.^9,49^ In addition, some groups that participated in round 41 included a term in their scoring function to encourage individual arabinose units to remain in a parallel stacking orientation with nearby aromatic residues in the active site. This type of protein–carbohydrate interaction, known as a CH–π interaction, has been well characterized in carbohydrate-binding proteins and is understood to play an important role in carbohydrate binding and recognition.^50,51^ The Rosetta software suite does not currently employ a scoring term to encourage this type of geometry-driven intermolecular interaction, and it might help further discriminate native-like oligosaccharide–protein interactions.

With fourteen successful predictions and two additional scoring successes, our performance in the rounds evaluated thus far was commensurate with other leading groups in the rounds we participated. Of our nine docking failures, we believe, retrospectively, that we had the sampling techniques available in Rosetta to better model targets 113, 126, 131, and 132. On the other hand, targets 114, 116, 117, 123, and 124 required blind prediction tools that do not yet exist and as a result, they did not elicit a successful model from any predictor. Broadly, the challenges that caused the most failures were docking with large conformational changes and multi-body docking (especially higher order heteromers). These community-wide failures highlight the massive gaps that still need to be addressed to fulfill the overarching goal of reliably modeling entire interactomes.^52–54^

## Acknowledgments

This work has been supported by grants from the National Institutes of Health, USA, *viz*. grants R01-GM078221 (JJG, SSRB, MLN, JRJ, JWL), T32-GM008403 (MLN, JRJ), F32-CA189246 (JWL) and F31-GM123616 (JRJ) and the National Science Foundation, USA award 1507736. Computations in this study have been performed in part on the Maryland Advanced Research Computing Center (MARCC) Blue Crab cluster. The authors thank Prof. Jamie Spangler of Johns Hopkins University for her advice on target 122.

