## Supplementary for "Novel sampling strategies and a coarse-grained score function for docking homomers, flexible heteromers, and oligosaccharides using Rosetta in CAPRI Rounds 37–45"

### Supplementary Information

**Contents**


| Contents | Page |
|---|---|
| Fig S1: SymDock2 model of T118 | S2 |
| Fig S2: Docking failures in Round 37 | S3 |
| Fig S3: Docking failures in Rounds 39 and 42 | S4 |
| Fig S4: Post-hoc analysis of T130 | S5 |

A

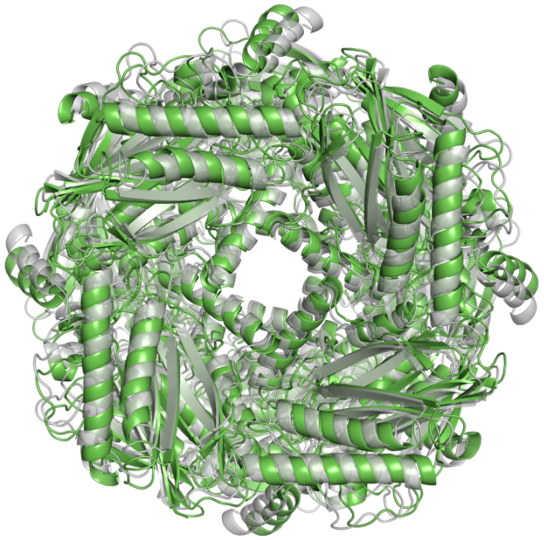

B

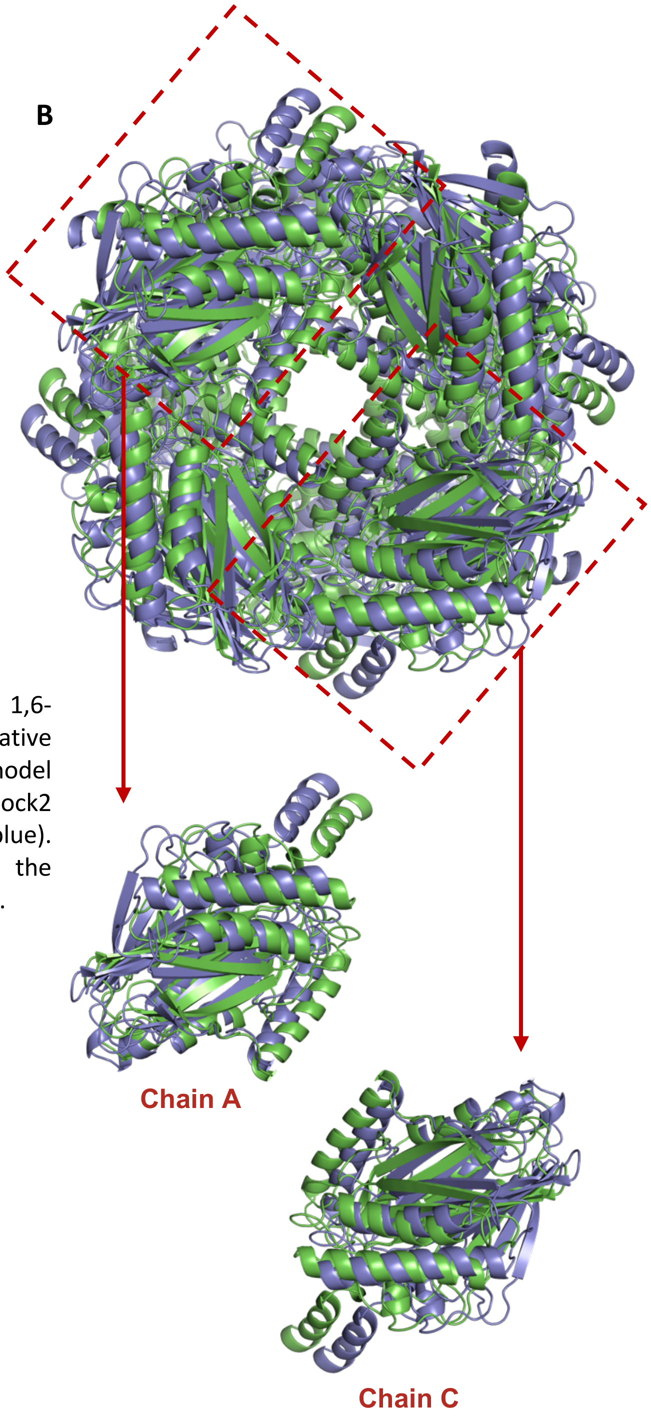


**Fig S1: T118** (A) Rosetta SymDock2 model of fructose 1,6-bisphosphatase (green) superimposed on the native approximation (gray, superposition of the monomer model to the crystal structure of a close homolog). (B) SymDock2 model (green) superimposed on the SymDock model (blue). Chains A and C are shown separately to highlight the increased distance in SymDock compared to SymDock2.

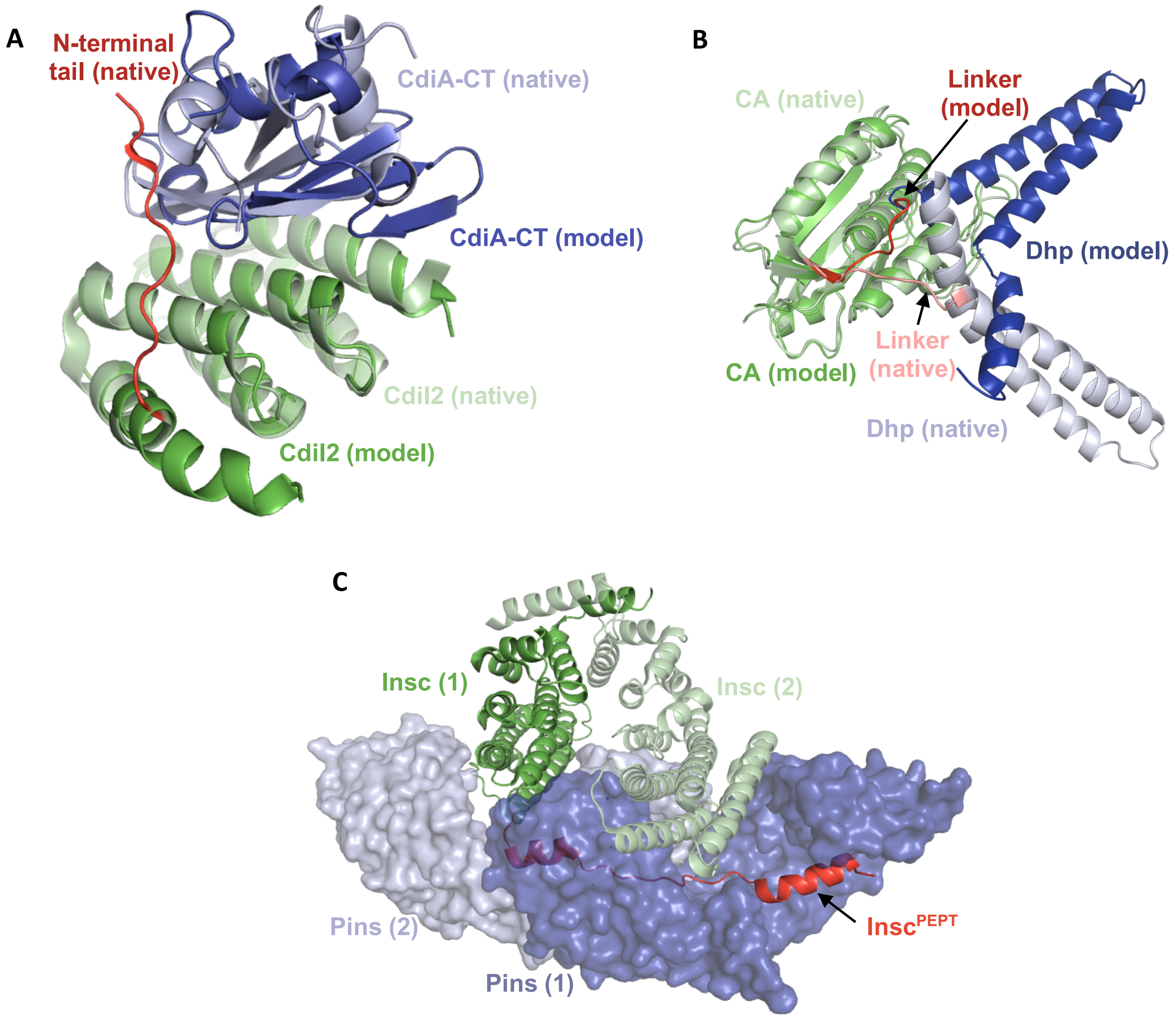


**Fig S2:** (A) **T113**: Model of CdiA-CT (blue) and CdiI2 (green) superimposed on the crystal structure of CdiI2 (pale green). The N-terminal region of CdiI2 was incorrectly predicted to be helical. Contacts with the N-terminal (red) are required for the proper rigid body orientation of CdiA-CT (pale blue). (B) **T116**: Monomer model CckA with the CA domain (pale green) superimposed on to that of the crystal structure (pale green). The incorrect conformation of the linker region in the model (red) displaces the Dhp domain (blue). The correct linker conformation (salmon) is required for the correct placement of Dhp (pale blue), and eventually the dimer. (C) **T117**: Crystal structure of the asymmetric Pins/Insc tetramer where the N-terminal region of Insc (red) binds the cavity inside Pins (blue). The rest of the Insc (green) does not show a strong interaction.

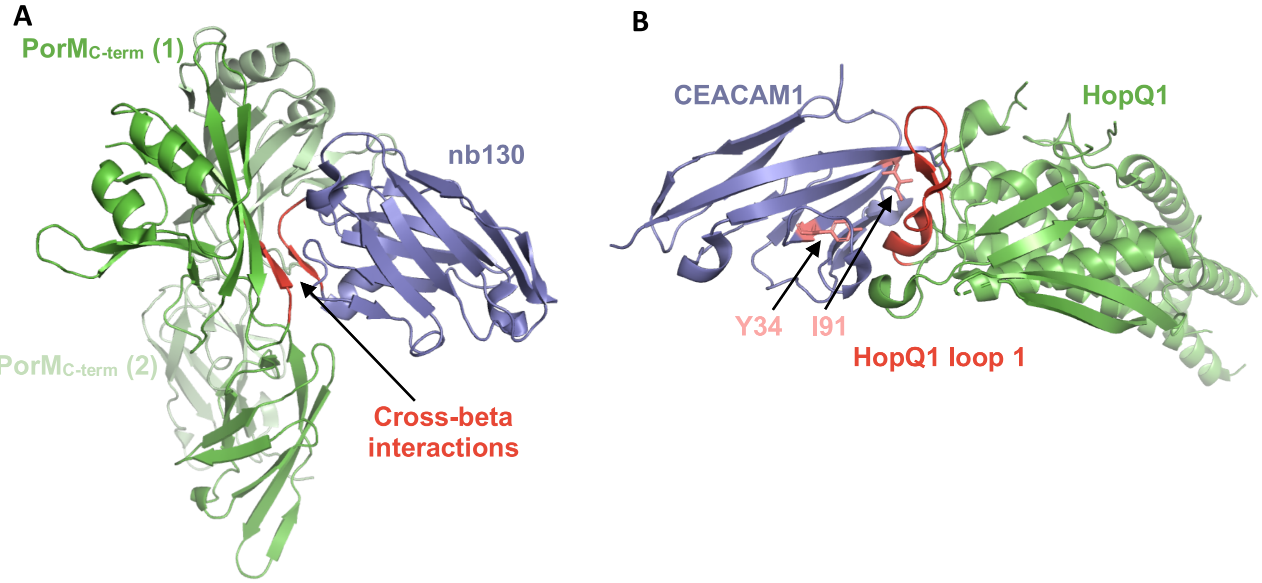


**Fig S3:** (A) **T123:** Structure of $PorM_{C\text{-}term}$ dimer (green) crystallized with the help of nb130 (blue). One nb130 molecule forms a cross-beta sheet at the interface with one of the $PorM_{C\text{-}term}$ molecules, thus helping them stabilize the interface and crystallize. (B) **T131**: Crystal structure of HopQ1 (green) bound to CEACAM1 (blue). The 14-residue loop of HopQ1 (red) is stabilized by interactions with CEACAM1, including the two residues predicted to be at the interface (salmon). The strand-turn-helix conformation adopted by the loop was not predicted correctly leading to incorrect rigid-body placement of CEACAM1.

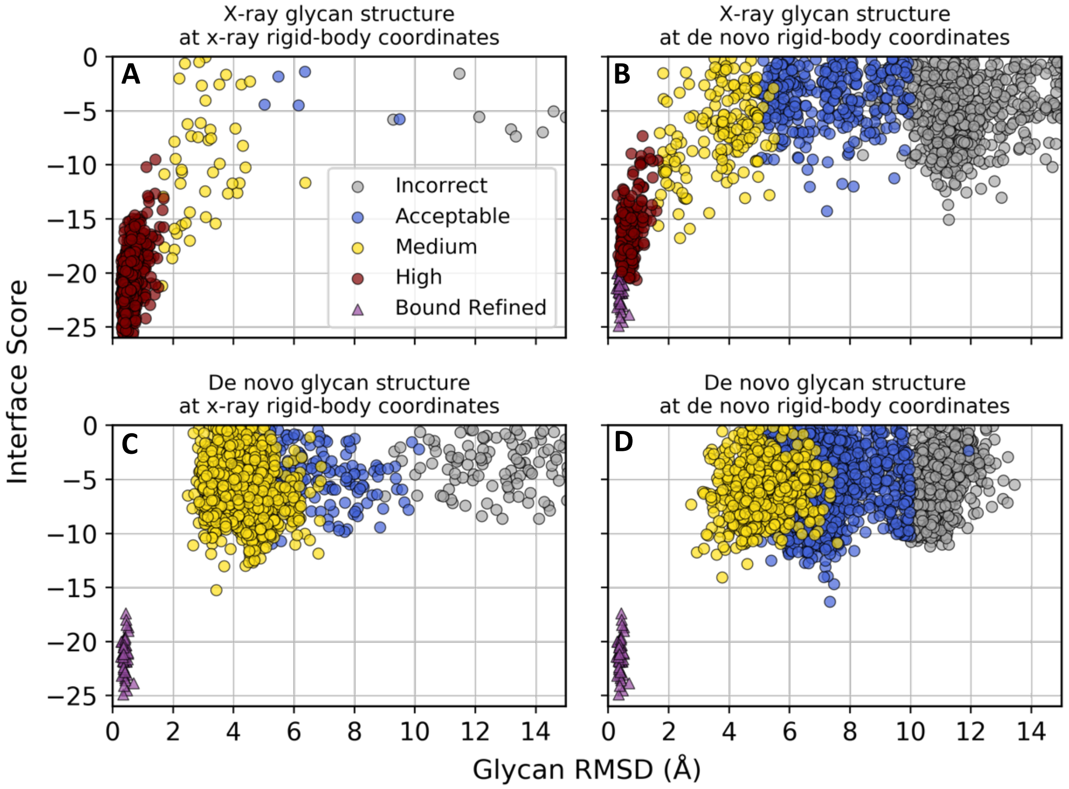


**Fig S4:** Glycan conformations generated using the GlycanDock algorithm on (A) the x-ray glycan structure, (B) the x-ray glycan structure aligned on each of the glycans of the three starting structures, (C) the glycans of the three starting structures each aligned on the x-ray glycan structure, and (D) the three starting structures employed during round 41. The bound refined data (purple triangles) represent glycan conformations generated by refining the native crystal structure, 6F1G. Colored circles indicate the CAPRI quality rating of each generated structure (1000 models each), all with the same protein structure used during round 41. All data were generated using the most current version of the GlycanDock algorithm at the time of writing. GlycanDock begins by performing an initial perturbation of ± 0.5 Å translation, ± 7.5° rotation, and backbone torsion perturbation between ± 12.5° on the input starting structure. This initial perturbation is then followed by a Monte Carlo simulation with ± 0.5 Å translations, ± 7.5° rotations, and backbone torsion moves between ± 30°. The weights of the attractive and repulsive terms of the Rosetta scoring function are set high and low and ramped slowly back down and up throughout ten cycles of Monte Carlo sampling, respectively, until reaching their standard weights in the last sampling cycle. Note that the bound refined simulations (purple triangles) do not perform the initial perturbation step, and only one simulation generating 50 decoys was performed with the same data being plotted in plots A–D for reference.
